# The neurotrophic factor artemin and its receptor GFRα3 mediate migraine-like pain via the ion channel TRPM8

**DOI:** 10.1101/2024.09.09.611532

**Authors:** Chenyu Yang, Chao Wei, Sanaa Alam, Xunyang Chen, David D. McKemy

## Abstract

**Background:** Migraine has a strong genetic foundation, including both monogenic and polygenic types. The former are rare, with most migraine considered polygenic, supported by genome-wide association studies (GWAS) identifying numerous genetic variants associated with migraine risk. Surprisingly, some of the most common mutations are associated with TRPM8, a non-selective cation channel that is the primary sensor of cold temperatures in primary afferent neurons of the somatosensory system. However, it is unlikely that the temperature sensitivity of TRPM8 underlies its role in migraine pathogenesis. To define the basis of the channel’s involvement, we reasoned that cellular processes that increase cold sensitivity in the skin, such as the neurotrophic factor artemin, via its receptor GFRα3, also mediate TRPM8-associated migraine-like pain in the meninges.

**Methods:** To investigate the role of artemin and GFRα3 in preclinical rodent migraine models, we infused nitroglycerin acutely and chronically, and measured changes in periorbital and hind paw mechanical sensitivity in male and female mice lacking GFRα3, after neutralization of free artemin with specific monoclonal antibodies, or by systemic treatment with a TRPM8-specific antagonist. Further, in wildtypes and mice lacking either GFRα3 or TRPM8, we tested the effects of supradural infusions of a mix of inflammatory mediators, artemin, and a TRPM8-specific agonist on migraine-related pain in mice.

**Results:** We find that mechanical allodynia induced by systemic nitroglycerin, or supradural infusion of inflammatory mediators, involves GFRα3. In addition, neutralization of circulating artemin reduces the nitroglycerin phenotype, demonstrating the importance of this neurotrophic pathway. Further, we show TRPM8 expression in the meninges and that direct supradural infusion of either a TRPM8-specific agonist or artemin itself produces mechanical allodynia, the latter dependent on TRPM8 and ameliorated by concurrent treatment with sumatriptan.

**Conclusions:** These results indicate that neuroinflammatory events in the meninges can produce migraine-like pain in mice via artemin and GFRα3, likely acting upstream of TRPM8, providing a novel pathway that may contribute to migraine pathogenesis.

## INTRODUCTION

Migraine is a strongly heritable neurovascular disorder, yet little is known of the underlying genetic basis for common migraine ^1, 2^. To this end, multiple genome-wide association studies (GWAS) have searched for single-nucleotide polymorphisms (SNPs) linked to either protection from migraine or associated with increased migraine susceptibility ^3–7^. Surprisingly, some of the most common mutations reside in or near the genetic locus of the menthol receptor, *transient receptor potential melastatin 8* (*trpm8*) ^3–8^, a cold-gated ion channel highly expressed in trigeminal ganglia (TG) ^9, 10^. While its role in cold somatosensation and nociception in the skin is well-studied ^11^, how TRPM8 channels might contribute to migraine pathogenesis is unclear ^12^.

The SNPs identified in *trpm8* reside in noncoding regions, suggesting they do not alter function but may influence channel expression ^5, 7, 13–20^. For example, the reduced migraine risk allele rs10166942[C] is located 5′ of the start codon ^3, 5^ and homozygous individuals express less TRPM8 and are less sensitive to cold than those with a T at this position ^21^. Intriguingly, this T allele, which is prevalent in individuals from higher latitudes and proposed to arise from positive selection for a thermoregulatory advantage in colder climates ^22^, is associated with increased migraine predisposition ^23, 24^. Thus, increased TRPM8 expression may correlate with increased migraine risk. What is also intriguing is that cold triggers headaches in humans (headache-induced by cold stimuli (HICS)), suggesting a link between headaches and TRPM8 activity ^25^.

Given its relative novelty as a migraine susceptibility gene, only a few preclinical studies have examined TRPM8 function related to migraine ^26–30^. Previously, using both mouse genetics and TRPM8-specific pharmacology, we showed that TRPM8 is required for both chronic and acute migraine-like phenotypes induced by systemic application of either nitroglycerin (NTG) or calcitonin gene-related peptide (CGRP) ^31^. However, the molecular mechanisms whereby TRPM8 channels and cells mediate migraine-like pain are not fully understood. To investigate this, we reasoned that processes underlying TRPM8-mediated cold allodynia and hyperalgesia in the skin could also facilitate the channel’s sensitization in migraine-relevant tissues. Specifically, the <u>g</u>lial cell-line <u>d</u>erived <u>n</u>eurotrophic factor (GDNF) artemin and its receptor GFRα3 are required for TRPM8-dependent cold allodynia and hyperalgesia in multiple pain models ^32–36^. Further, artemin expression is present in meningeal vascular smooth muscle and GFRα3 expressing nerve fibers are found throughout the dura mater ^37, 38^. Expression of both meningeal GFRα3 and vascular artemin in the rat increases after nitroglycerin (NTG) treatment ^38^, and artemin is proposed to increase expression of the inducible form of nitric oxide synthase (iNOS) in trigeminal neurons ^39^. Thus, there is circumstantial evidence for a role of artemin/GFRα3 signaling in the meninges that may correlate with migraine pathogenesis.

Here, we report that, like TRPM8, GFRα3 is required for NTG-induced acute and chronic tactile allodynia, a phenotype that is also attenuated by neutralization of artemin. Further, GFRα3 is involved in allodynia produced by a supradural infusion of a mixture of inflammatory mediators. Lastly, application of either a TRPM8 agonist or artemin directly onto the mouse dura matter leads to similar phenotypes that are TRPM8-dependent, as well as sensitive to the abortive migraine agent sumatriptan. Our findings suggest a conserved mechanism underlies both TRPM8-dependent cold allodynia and migraine-like pain, highlighting a novel molecular pathway that may, in part, contribute to migraine headaches.

## MATERIALS & METHODS

### Animals

All experiments were approved by University of Southern California Institutional Animal Care and Use Committee and in compliance with ARRIVE guidelines for Animal Research: Reporting of In Vivo Experiments ^40^. Mice aged from 8 weeks to 16 weeks were used in all experiments and unless otherwise specified, equal numbers of male and female mice were used. All mice were housed in a temperature-controlled environment on a 12-hour light and dark cycle with ad libitum access to food and water. All experiments were performed during the light cycle from 8 AM to 5 PM. Trpm8-null mice (Trpm8^-/-^, Strain # 008198) ^41^ were purchased from Jackson Laboratories and were in the C57Bl/6 background. Wildtype C57Bl/6 and Trpm8^GFP^ ^10^ mice were maintained in-house, as were Gfrα3-null (Gfrα3^-/-^) mice ^42^ that were bred from Gfrα3^+/-^ animals to obtain Gfrα3-nulls and wildtype littermate controls, as described ^34^.

### Supradural Injections

Supradural injections were performed as previously described ^43^. Briefly, mice were anesthetized with 3%-4% isoflurane and then maintained by 1.5%-2% isoflurane through a nose cone. The various agents used were injected in a volume of 5 µl using an internal cannula (Invivo1, part #C313IS-5/SPC, Internal Canula, 28 gauge) connected to plastic tubing (Perkin Elmer, Cat#N0695476, 2-stop, I.D. 0.19 mm) and a 10 µl glass syringe (Hamilton, Cat# 80300). The inner projection of the cannula was adjusted to be 0.6 mm to inject through the soft tissue at the intersection of lambdoidal and sagittal sutures but not puncture the dura. Post-mortem analysis was done to confirm that the dura remained intact after each injection.

### Retrograde Labeling of Trigeminal Neurons Innervating Mouse Dura

Trpm8^GFP^ mice were anesthetized and 5 μl supradural injections of the retrograde tracer wheat germ agglutinin (1 mg/ml, WGA-Alexa-350; Cat#W11263, ThermoFisher) were performed as described ^43^. After recovery from anesthesia, mice were housed in their cages for 14 days to allow for the transport of the tracers to the dural afferent somata in the trigeminal ganglia (TG).

### Tissue Preparation and Immunohistochemistry

TG were dissected from Trpm8^GFP^ mice injected with WGA tracer as previously described^10^. Briefly, the mice were euthanized and perfused transcardially with Phosphate Buffer Solution (PBS) followed by fixative (cold 4% paraformaldehyde solution, PFA). Tissues were carefully removed and post-fixed in the same fixative for another 2 hours at 4°C. After cryoprotection in 30% sucrose solution for at least 24 hours, the tissues were fast-frozen in OCT and sectioned with a cryostat at 16 µm. The tissue sections were mounted on Superfrost Plus slides and kept at -80°C. On the day of immunostaining, frozen slides were dried at 25°C for 30 mins, washed with deionized water and PBS for three times, and permeabilized in 0.1% Triton + PBS (PBST) for 30 mins. The mounted sections were blocked in 0.1% PBST solution with 20% Normal Goat Serum (NGS) for 1 hour. The slides were incubated at 4°C overnight with primary antibodies in 0.1% PBST solution with 0.5% BSA + 0.5% NGS (Chicken anti-GFP antibody, 1:500, Cat# GFP-1020, Aves Lab; Rabbit anti-CGRP antibody, 1:1000, Cat# C8198, ThermoFisher). The slides were washed in incubation buffer 6x5 mins and incubated in secondary antibodies for another 2 hours at room temperature (Goat anti-Chicken-IgY antibody, Alexa-488, 1:2000, Cat# A11039, ThermoFisher; Goat anti-Rabbit-IgG antibody, CF594, 1:2000, Cat# SAB4600234, Sigma Aldrich). The slides were washed in incubation buffer 6x5 mins followed by 2x5 mins PBS and covered by media and coverslips. The slides were dried and kept at 4°C overnight before imaging.

The meninges (including dura mater) were dissected from Trpm8^GFP^ mice using a protocol adapted from previous reports ^44^. Briefly, mice were euthanized and perfused transcardially with ice-cold PBS. Skulls with meninges attached were removed and fixed in 2% PFA solution at 4°C overnight. The meninges were carefully detached from skulls in PBS solution the next day and transferred to a 24-well plate. The tissues were blocked for 1 hour in 0.1% PBST solution with 2% NGS + 1% BSA and then incubated overnight at 4°C in 0.1% PBST solution with 1% BSA and primary antibodies. After 3x5 mins wash with PBS, the tissues were incubated in 0.1% PBST solution with 1% BSA and secondary antibodies at room temperature for 2 hours. The tissues, after 3x5 mins wash with PBS, were flattened and dried in the dark on Superfrost Slides for 10-15 mins, covered with appropriate media and coverslips and imaged immediately.

### Image Acquisition and Analysis

Digital images were acquired using a Zeiss Axio Imager.M2 (Oberkochen, Germany) with Apotome attachment and Zeiss Zen Blue software. Quantification of overlap between expression of neuronal markers, retrograde tracers, and GFP was obtained per field and expressed as percentage overlap with the mean ± standard error of the mean between fields.

### Pharmacological agents and migraine-like pain models

Nitroglycerin (NTG) was purchased from VWR (Cat# 101095-632) and supplied as 1% solution in Propylene Glycol (PG) and further diluted to the desired working concentration (1 mg/kg or 10 mg/kg) using a 0.9% saline solution. NTG or vehicle was administered intraperitoneally (i.p.) every other day for 9 days, as described ^31, 45^, with mechanical sensitivity measured before and 2 hours after each injection. Icilin was purchased from Tocris Bioscience (Cat# 1531), dissolved, and diluted to 1 nmol / 5 µl in PG and injected supradurally as described above. PBMC was purchased from Focus Biomolecules (Cat# 10-1413), dissolved, and diluted to working concentration (4 mg/kg) using 10% DMSO in a 0.9% saline solution and injected i.p. as described ^32, 46^. Artemin was purchased from R&D Systems (Cat# 1085-AR), dissolved, and diluted to 1 pg / 5 µl using a Hank’s Balances Salt Solution (HBSS: 140 mM NaCl, 5 mM KCl, 1 mM CaCl_2_, 0.4 mM MgSO_4_, 0.5 mM MgCl_2_, 0.3 mM Na_2_HPO_4_, 0.4 mM KH_2_PO_4_, 6 mM D-glucose, 4 mM NaHCO_3_) and injected supradurally. The inflammatory soup (IS) components were purchased from Sigma Aldrich, consisting of 1 mM Bradykinin (Cat# 05-23-0500), 1 mM 5-HT (Cat# H9523), 1 mM Histamine (Cat# 59964), 200 µM PGE_2_ (Cat# P5640) and at pH 5.0 in Synthetic Interstitial Fluid (SIF: 10 mM HEPES, 5 mM KCl, 135 mM NaCl, 1 mM MgCl_2_, 2 mM CaCl_2_, 10 mM glucose). Vehicle solutions were used as control injections in experiments accordingly. The IS or vehicle was injected once supradurally and mechanical sensitivity was measured from one to six hours post-injection, then again twenty-four hours later. The anti-artemin monoclonal and control IgG Isotype monoclonal antibodies were purchased from R&D Systems (Artemin: MAB1085, Control: MAB002) and reconstituted in sterile PBS solution. The antibodies were injected intraperitoneally at a concentration of 10 mg/kg. Sumatriptan was purchased from Sigma Aldrich (Cat# S1198), diluted using 0.9% Saline solution, and injected subcutaneously (s.c.) at a concentration of 0.6 mg/kg.

### von Frey Assay

The von Frey assay was used to assess the sensitivity to mechanical stimulation as previously described ^31, 43^. Mechanical sensitivity (50% withdrawal threshold) was determined at the timepoints indicated with calibrated von Frey monofilaments (Semmes-Weinstein) using the up-and-down method. Briefly, the mice tested were conditioned by handling for 5 minutes and habituated to plexiglass chambers on metal mesh (hindpaw) or horizontal paper cups (periorbital assay) for 3 consecutive days before experiments. The filaments (for hind paw assay: starting from 0.4 g filament; flexible force range of 0.008-2 g; for periorbital assay: starting from 0.07 g filament, flexible force range of 0.008g-0.6 g) were applied to the plantar surface of the hind paw or the center of the face between eyes for up to 5-6 seconds until the filament bent or positive response was recorded. The next heaviest (if no response) or lightest (if a response) filament was used in the next measurement and 4 additional measurements were performed. The 50% withdrawal threshold was calculated using the SUDO method previously described ^47^. The mice were returned to their home cage after measurements and retested on the next treatment/test day.

### Statistical Analysis

The experimenters were blinded to either genotype of the mice receiving treatments or substances injected. Unless otherwise specified, eight to ten mice were tested in each group and equally balanced between male and female mice. No sex differences were observed in the data. All data are presented as mean ± standard error of the mean (SEM) with the statistical assay used described in the figure legend or associated text, along with sample numbers. Time course measurements and comparisons between different injection conditions such as treatment or genotype were made with were determined via a two-way ANOVA with either Bonferroni’s, Dunnett’s, or Tukey’s multiple comparison tests. Two-tailed paired t-tests were used to compare pre-versus post-injection conditions with the same injection conditions or genotype.

## RESULTS

### TRPM8 afferents are present in the mouse dura

To further investigate the role of TRPM8 in migraine-like pain, we confirmed that TRPM8-positive afferents are present in the mouse meninges, first using retrograde tracer back-labeling ^43^. Naïve adult Trpm8^GFP^ mice ^10^ received a supradural injection of the retrograde tracer wheat germ agglutinin (1%; WGA-Alexa-dye) and TG were processed for immunolabeling 14 days post-injection as described ^10, 34, 35, 48^. We found that 15.7±2.7% of WGA-labeled cells were GFP positive (Fig. 1A, n=4 mice; 805 WGA-positive cells), consistent with the proportion of TG neurons per ganglia known to express TRPM8 ^9, 10, 34, 49, 50^. Next, we performed immunolabeling of dura dissected from adult Trpm8^GFP^ mice as described ^10, 34, 51^, observing GFP expression in Trpm8^GFP^ animals (Fig. 1B), labeling that was absent in wildtype animals (Fig. 1C). These results are consistent with prior studies showing TRPM8 expressed in this tissue ^52^. Of note, a proportion of GFP-positive axons also showed immunoreactivity for CGRP, consistent with soma co-expression of TRPM8 and CGRP in the trigeminal ganglia ^9, 10, 34, 49, 50^.

**Figure 1:**
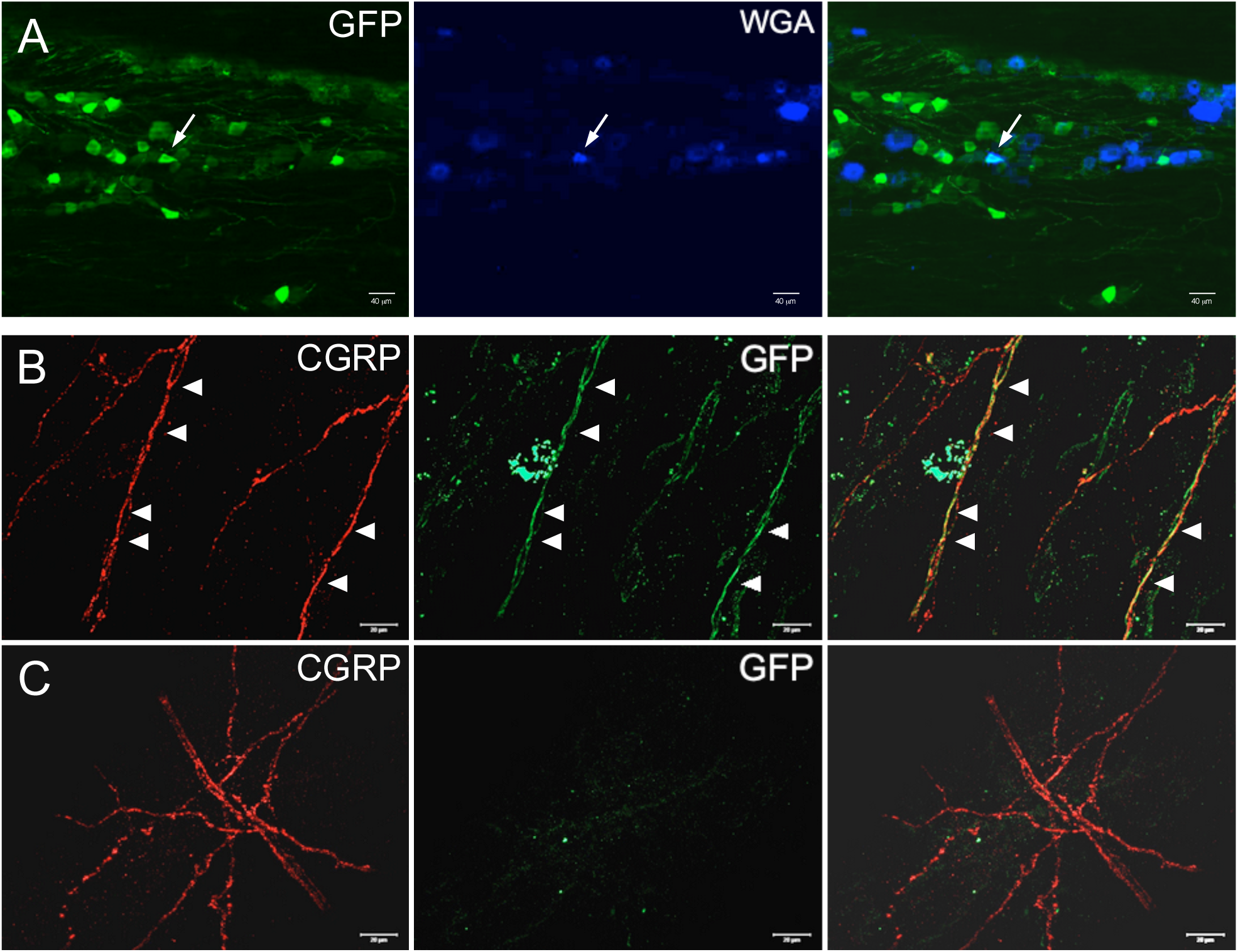
TRPM8-positive afferents innervate mouse meninges. (A) Representative images of GFP (TRPM8) and WGA (Alex-350) co-labeling in trigeminal ganglia in adult Trpm8^GFP^ mice that received an intradural injection of WGA. Arrows highlight co-labeled cells. Immunoreactivity for GFP (TRPM8) and calcitonin gene-related peptide (CGRP) in the meninges of Trpm8^GFP^ mice (B) versus that in wildtype mice (C). Arrowheads denote axons that co-label for GFP and CGRP.

### GFRα3 is required for nitroglycerin-induced tactile allodynia

Next, we asked how activation of TRPM8 channels in the meninges, which are likely not responding to temperature in this tissue, might lead to migraine-like pain. We reasoned that mechanisms known to induce TRPM8-dependent cold allodynia in the skin might also be relevant in the dura matter. The neurotrophic factor artemin, by binding to its cognate receptor GDNF Family Receptor α3 (GFRα3), can induce TRPM8-dependent cold hypersensitivity, and GFRα3 is also required for or involved in inflammatory and neuropathic cold pain of various etiologies, including migraine ^32–35^. To test this premise, we measured acute periorbital mechanical allodynia in wildtype and GFRα3-null mice (Gfrα3^-/-^) ^34, 42^ 2 hours after they were given an intraperitoneal injection (i.p.) of 1 mg/kg NTG or vehicle, repeated every 2 days ^31, 45^ (Fig. 2A). As expected, wildtype animals exhibited mechanical allodynia after each NTG injection that was significantly different than baseline (BL) at all days tested, as well as compared to vehicle injected mice by the 2^nd^ injection on Day 3 (p<0.01, 2-way ANOVA with Tukey’s multiple comparisons test). In contrast, Gfrα3^-/-^ mice were insensitive to NTG treatment as their mechanical sensitivity was not different than baseline or compared to vehicle injected animals (p>0.05, 2-way ANOVA with Tukey’s multiple comparisons test) at all days tested. To ensure that the lack of a phenotype in Gfrα3^-/-^ mice wasn’t due to an NTG dosage effect, we also tested their acute response to 10 mg/kg NTG, again finding that the robust increase in mechanical allodynia in wildtype mice 2 hours after a single injection was absent in Gfrα3^-/-^ animals (Fig. 2B). Lastly, to determine if GFRα3 was also required for extracephalic allodynia induced by NTG, we measured hind paw mechanical responses in wildtype and Gfrα3^-/-^ mice before and 2 hrs after an i.p. injection of 10 mg/kg NTG, observing no differences in Gfrα3^-/-^ mice given NTG (Fig. 2C).

**Figure 2:**
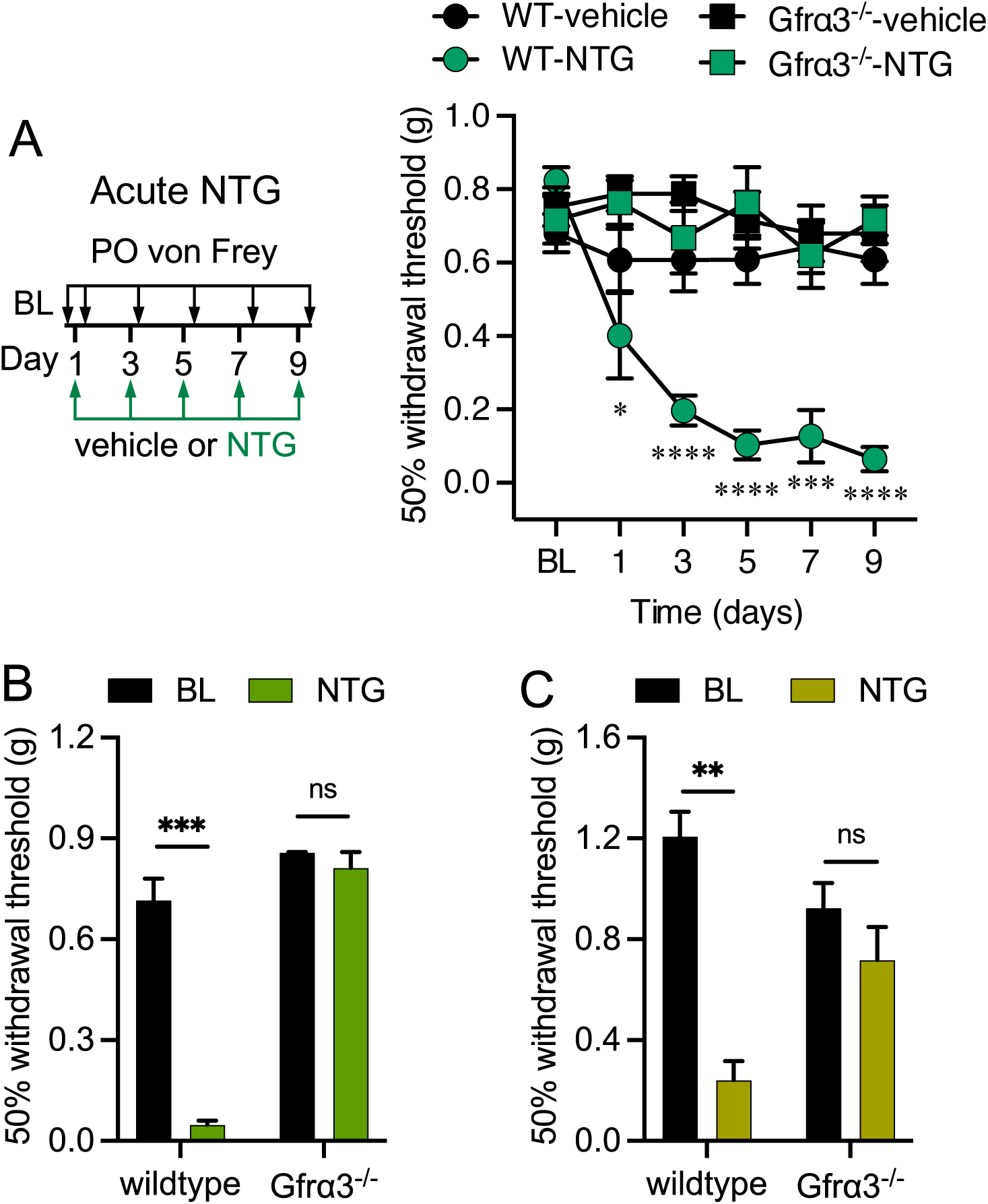
GFRα3 is required for nitroglycerin-induced acute cephalic and extracephalic tactile allodynia. (A) Left: Schematic of acute injections and mechanical testing timeline. Right: Wildtype mice exhibit periorbital mechanical allodynia measured 2 hrs after each systemic NTG injection (1 mg/kg) compared to baseline (BL) sensitivity (n=8 for each condition, two-way ANOVA with Dunnett’s multiple comparison test, *p<0.05, ***p<0.001, ****p<0.0001). In contrast, GFRα3-null mice (Gfrα3^-/-^) are insensitive to 1 mg/kg NTG injections on each day tested (n=6 for each condition, two-way ANOVA with Dunnett’s multiple comparison test compared to BL, p>0.05). Wildtypes receiving a 10 mg/kg dose of NTG show robust periorbital (B) and hindpaw (C) mechanical allodynia 2 hr post-injection compared to baseline (BL), a phenotype absent in Gfrα3^-/-^ mice (n=6 and 8 for periorbital and hindpaw, respectively, paired two-tailed t-test, ^ns^p>0.05, **p<0.01, ***p<0.001).

Repeated NTG dosing leads to an increase in basal mechanical sensitivity that is considered a potential model of chronic migraine ^31, 45^. To test the role of GFRα3 in chronic migraine, we compared baseline behaviors at each injection day prior to treatment with 1 mg/kg NTG or vehicle, observing the expected increase in basal allodynia in wildtypes, an effect that was absent in Gfrα3^-/-^ animals (Fig. 3A). We again assessed 10 mg/kg NTG, also observing that basal mechanical allodynia was absent in Gfrα3^-/-^ mice 2 days after a single NTG injection (Day 3, Fig. 3B). Similarly, hindpaw mechanical allodynia did not develop in Gfrα3^-/-^ mice given 10 mg/kg NTG (Fig. 3C). Thus, GFRα3 is required for both cephalic and extracephalic tactile allodynia in both acute and chronic models of rodent migraine-like pain, a phenotype like that observed in TRPM8-null mice ^31^.

**Figure 3:**
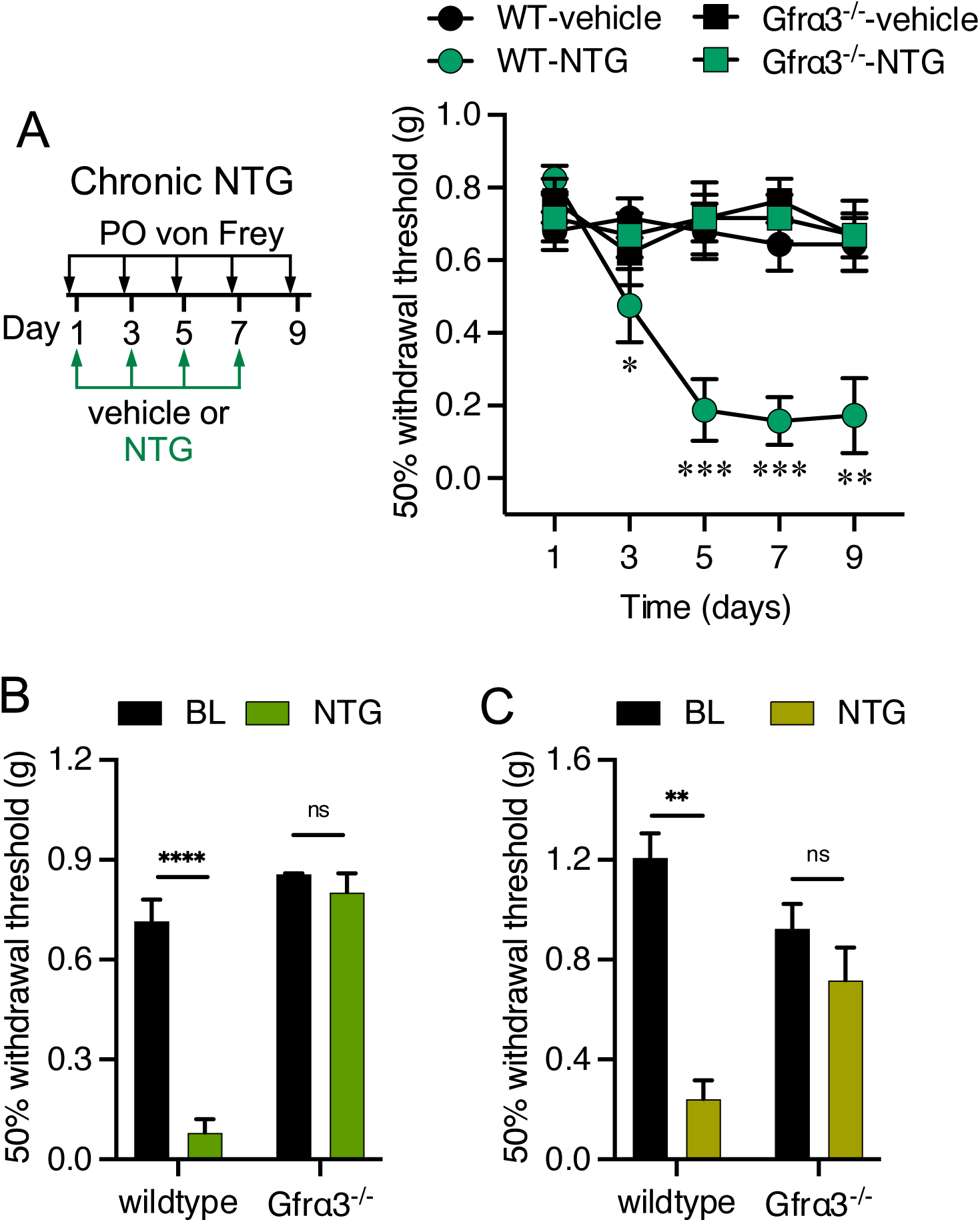
GFRα3 is required for chronic allodynia induced by repeated nitroglycerin treatments. (A) Left: Schematic of chronic injections and mechanical testing timeline. Right: Wildtype and Gfrα3^-/-^ mice received 1 mg/kg NTG injections every two days, with only wildtypes displaying enhanced basal periorbital mechanical allodynia, compared to baseline at Day 1 (n=6-8 for each condition, two-way ANOVA with Dunnett’s multiple comparison test, *p<0.05, **p<0.01, ***p<0.001). Wildtypes receiving a 10 mg/kg dose of NTG show robust basal periorbital (B) and hindpaw (C) mechanical allodynia 2 days post-injection compared to baseline (BL), a phenotype is absent in Gfrα3^-/-^ mice (n=6 each condition for periorbital and hindpaw, respectively, paired two-tailed t-test, ^ns^p>0.05, **p<0.01, ****p<0.0001).

### Neutralizing artemin reduces nitroglycerin-induced tactile allodynia

Our results show that GFRα3 is required for NTG-induced tactile allodynia, suggesting that its specific ligand artemin might also play a role in this phenotype. To test this, we employed monoclonal antibodies (mAb) raised against artemin that we have shown can specifically ameliorate pathological cold allodynia. ^32–34, 53^. Wildtypes were tested for basal cephalic mechanical sensitivity then injected with either an anti-artemin mAb or an isotype control. The mice were re-tested at 1 hour post-injection, demonstrating that mAb treatment did not alter basal mechanosensation (Fig. 4A). Immediately after testing, mice treated with the two mAbs were injected with 1 mg/kg NTG and re-tested 2 hours post-injection. Mice given the isotype control showed robust mechanical allodynia, as expected, whereas mice injected with the anti-artemin mAb exhibited normal sensitivity (Fig. 4A).

**Figure 4:**
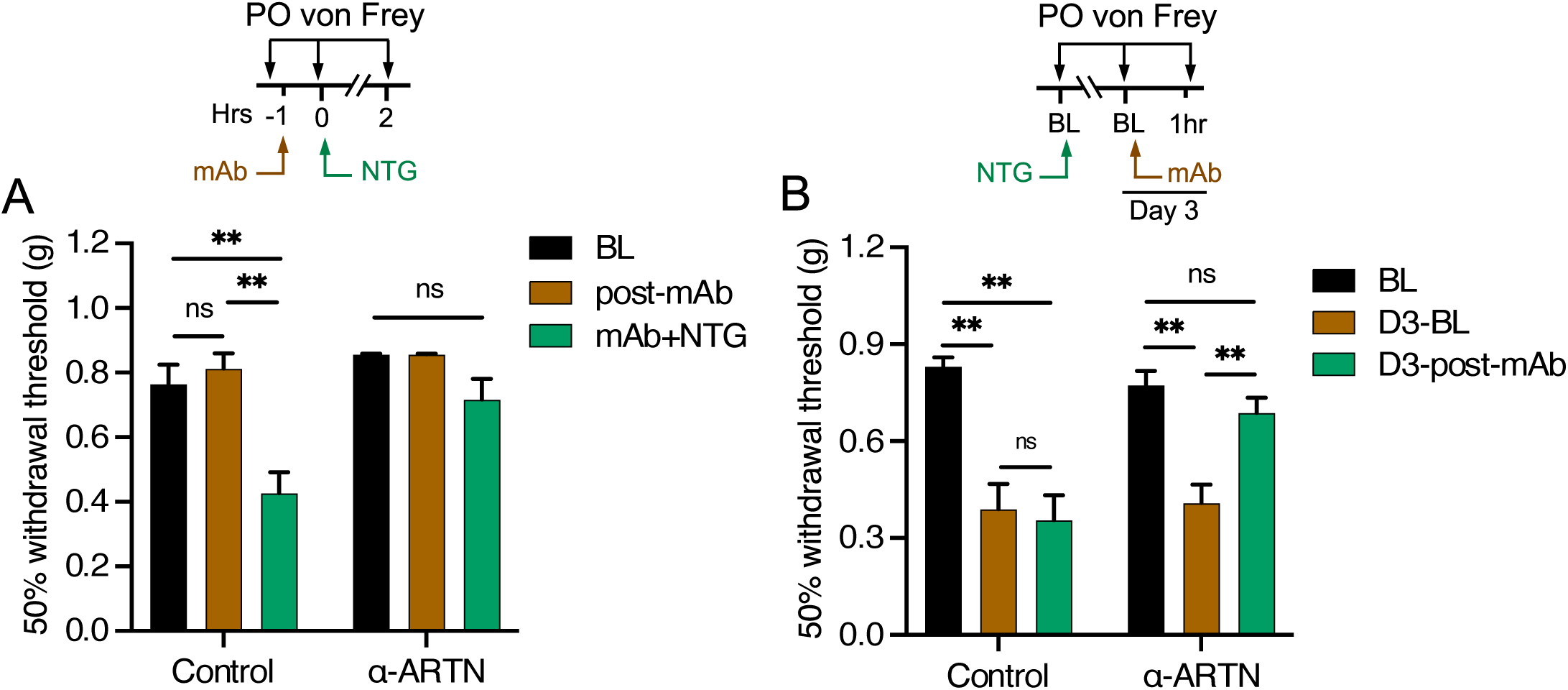
Neutralizing artemin ameliorates nitroglycerin-induced tactile allodynia. (A) Top: Schematic of the experimental approach. Wildtype mice treated with either an anti-artemin or isotype control mAb exhibit normal periorbital mechanical sensitivity 1hr post-injection compared to baseline (BL). A subsequent injection of 1 mg/kg NTG induced allodynia in mice given the isotype control, but not the anti-artemin mAb (n=6 for each condition, one-way ANOVA with Tukey’s multiple comparisons tests for each time point and condition, ^ns^p>0.05, **p<0.01). (B) Top: Schematic of the experimental approach. Wildtype mice treated with 1 mg/kg NTG to induce basal periorbital mechanical allodynia 2 days post-injection compared to BL, a phenotype that remained in mice subsequently treated with the isotype control mAb. However, allodynia was reduced in animals treated with the anti-artemin mAb (n=10 for each condition, one-way ANOVA with Tukey’s multiple comparisons test for different time point and condition, **p<0.01, ^ns^p>0.05).

Next, we asked if chronic allodynia induced by NTG was sensitive to artemin neutralization. Animals were tested for basal mechanical sensitivity (Day 1) and given a single injection of 1 mg/kg NTG. Their basal sensitivity was remeasured 2 days later (Day 3), with all animals exhibiting mechanical allodynia (Fig 4B). These mice were then given either an injection of the isotype control or anti-artemin mAbs and tested 1 hour later. As with acute sensitization, artemin neutralization decreased mechanical allodynia, bringing the 50% withdrawal threshold back to basal levels. These data indicate that both acute and the initial phase of chronic mechanical allodynia induced by NTG are dependent on free, circulating artemin, as well as suggest that established allodynia can be alleviated by artemin neutralization.

### Dura artemin treatment induces TRPM8-dependent tactile allodynia

Our results suggest a key role for artemin and GFRα3 signaling in the NTG model of migraine-like pain but doesn’t directly determine if GFRα3 is acting at sites involved in migraine, such as the meninges. To address this, we first asked if periorbital mechanical allodynia induced by stimulation of the meninges via direct supradural injection of a cocktail of inflammatory mediators required GFRα3 ^43^. Wildtype and Gfrα3^-/-^ mice were tested for basal (BL) mechanical sensitivity as above, then given a 5 μl injection of either vehicle (SIF; synthetic interstitial fluid) or an inflammatory soup (IS) mixture (in SIF at pH 5.0). Both wildtype and Gfrα3^-/-^ mice given IS showed robust mechanical allodynia, compared to vehicle injected mice (wildtypes: p<0.05 at 1 hour, p<0.0001 at 2-4 hours post-injection; Gfrα3^-/-^: p<0.01 at 2 hours, p<0.05 at 3hrs, two-way ANOVA with Tukey’s multiple comparison tests) and basal sensitivity (wildtypes: p<0.01 at 1 hour, p<0.0001 at 2 to 4 hours post-injection; Gfrα3^-/-^: p<0.01 at 2 hours, p<0.001 at 3 hours, p<0.05 at 4 hours post-injection; two ANOVA with Dunnett’s multiple comparison test), for several hours post-injection. However, the level of sensitization after IS injection was significantly lower in Gfrα3^-/-^ mice compared to wildtypes at 2-4hrs post-injection (Fig. 5A), indicating that GFRα3 mediates, in part, migraine-like pain produced by meningeal stimulation with inflammatory mediators.

**Figure 5:**
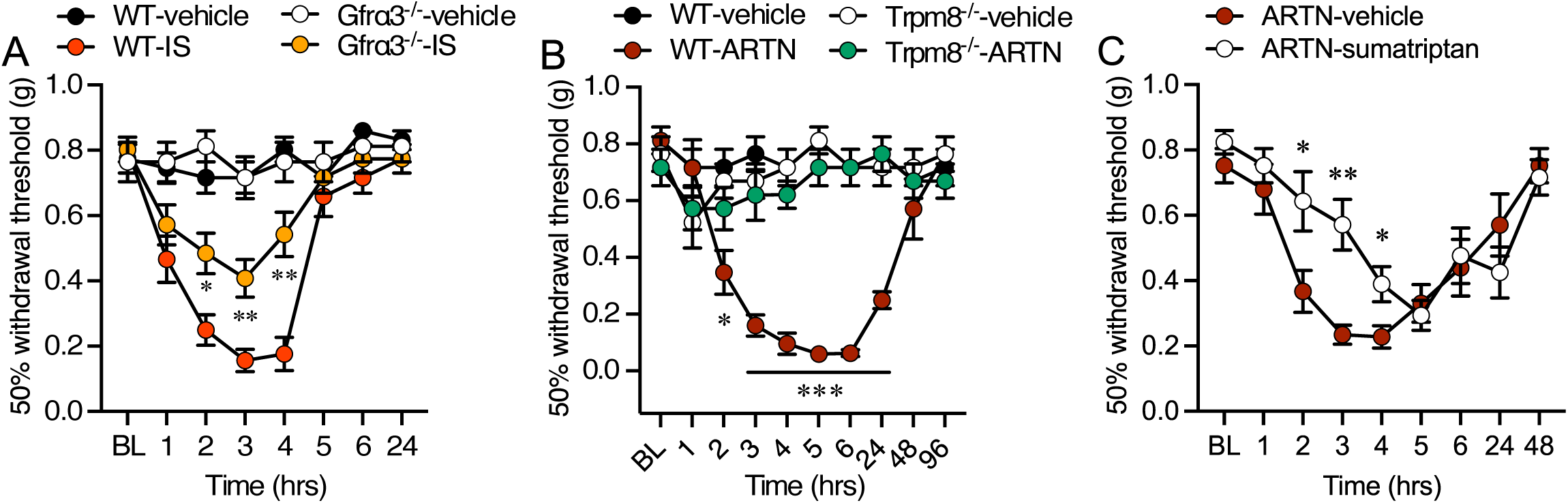
Necessity of GFRα3 and artemin in tactile allodynia induced by direct stimulation of the meninges. (A) Direct supradural infusion of an inflammatory soup (IS) mixture induces periorbital tactile allodynia that is partially dependent on GFRα3. Gfrα3^-/-^ mice given IS exhibited a reduced level of mechanical allodynia compared to wildtypes at 2-4hrs post-injection (n=10 each genotype, two-way ANOVA with Tukey’s multiple comparisons test, *p<0.05, **p<0.01). (B) Supradural artemin induces robust periorbital tactile allodynia in wildtype but not Trpm8^-/-^ mice (n=6 for each condition and genotype, two-way ANOVA with a Tukey’s multiple comparisons test for vehicle vs. artemin at each time point; *p<0.05, ***p<0.001). (C) Wildtype mice treated with artemin and sumatriptan exhibit a partial reduction in periorbital mechanical allodynia compared to those given artemin and vehicle (n=8 for each condition, *p<0.05, **p<0.01, two-way ANOVA with a Tukey’s multiple comparison test).

Next, we asked if direct application of artemin to the meninges in wildtype mice could induce a similar migraine-like phenotype. Naïve wildtype mice were tested for baseline periorbital mechanical sensitivity, given a supradural injection of either vehicle (HBSS) or 1 pg (5 μl) artemin, then retested for mechanical allodynia over the next 96 hours. Consistent with our Gfrα3^-/-^ and artemin neutralization results, we observed robust allodynia in mice receiving artemin that lasted over 24 hours post-injection (Fig. 5B). Thus, like other inflammatory agents, artemin directly induces tactile allodynia when applied to the dura. Next, as artemin induces TRPM8-dependent cold hypersensitivity in the skin ^32, 34, 35^, we reasoned that artemin and GFRα3 signaling underlies the TRPM8-dependent migraine-like phenotypes we observed ^31^. To test this hypothesis, we repeated the dura artemin injections in TRPM8-null mice (Trpm8^-/-^) ^41^, finding that this treatment did not induce mechanical allodynia in the absence of TRPM8 (Fig. 5B). Thus, as with cold allodynia in the skin, artemin-induced sensitization in the meninges is TRPM8 dependent. Lastly, to further define this artemin phenotype, we concurrently treated wildtype mice with supradural artemin and 0.6 mg/kg (subcutaneously) sumatriptan, the latter a traditional migraine treatment in humans and an agent used in animal studies to determine if a phenotype is migraine-like pain ^54–56^. Mice given artemin and the vehicle for sumatriptan (0.9% Saline) again showed robust mechanical allodynia starting at 2 hours post-injection (p<0.5 at 2 and 6 hours, p<0.0001 at 3 and 4 hours, p<0.01 at 5 hours post-injection, n=8, two-way ANOVA with Tukey’s multiple comparisons test). In contrast, mice given artemin and sumatriptan exhibited reduced mechanical allodynia from 2 to 4 hours post-injection, compared to artemin/vehicle treated mice (Fig. 5C), results suggesting that the artemin phenotype is consistent with migraine-like pain.

### Activation of meningeal TRPM8 induces cephalic tactile allodynia independent of GFRα3

As we find that artemin neutralization alleviates both acute and chronic tactile allodynia induced by NTG, and artemin-induced mechanical allodynia is TRPM8-depedent, we posit that this effect of artemin is via TRPM8 sensitization. Previously, we showed that acute extracephalic allodynia induced by 10 mg/kg NTG could be prevented by pre-treatment with the TRPM8 antagonist PBMC, but that established chronic NTG measured after 4 NTG treatments given every two days was insensitive to TRPM8 inhibition ^31^. Although inconsistent with our results in Fig. 4B, these prior results were with a substantially higher dose of NTG (10mg/kg versus 1mg/kg) given repeatedly. Thus, to determine if TRPM8 underlies the basal sensitization shown in Fig. 4B, wildtype mice were given a single 1 mg/kg NTG injection, then tested for basal allodynia 2 days later (BL-D3) with all animals exhibiting sensitization (n=13, paired two-tailed t-test, p<0.0001). These mice were randomly separated into two groups with one receiving an injection of 4 mg/kg PBMC and the other vehicle, then tested for mechanical sensitivity every hour for 5 hours. Mice given vehicle remained allodynic over this time period as there were no significant differences in mechanical sensitivity compared to the Day 3 baseline (D3-BL; p>0.05, two-way ANOVA with Dunnett’s multiple comparison test) and remained significantly different than the responses measured pretreatment (BL) at all time points tested (p<0.01 two-way ANOVA with Dunnett’s multiple comparison test). Conversely, mice given PBMC showed transient relief from this allodynia by 2 hours post-PBMC compared to the Day 3 baseline and to vehicle injected animals (Fig. 6A). Thus, inhibition of TRPM8 systemically relieved the initial NTG induced basal mechanical allodynia in this chronic migraine model. Further, to contrast with our prior results, mice were similarly given 4 injections of 1 mg/kg NTG at 2-day intervals, then their baseline (BL-D9) was measured 2 days after the 4^th^ injection (Day 9), with all animals exhibiting robust allodynia (Fig. 6A, paired two-tailed t-test, p<0.0001). In contrast to its effects on Day 3, mice given 4 mg/kg PBMC on Day 9 remained allodynic with periorbital mechanical allodynia that was like vehicle treated animals (Fig. 6A). These results are consistent with previous findings and demonstrate that, while the initial basal allodynia induced with 1 dose of NTG is TRPM8 and artemin dependent, the robust phenotype developed by repeated NTG treatments is independent of TRPM8.

**Figure 6:**
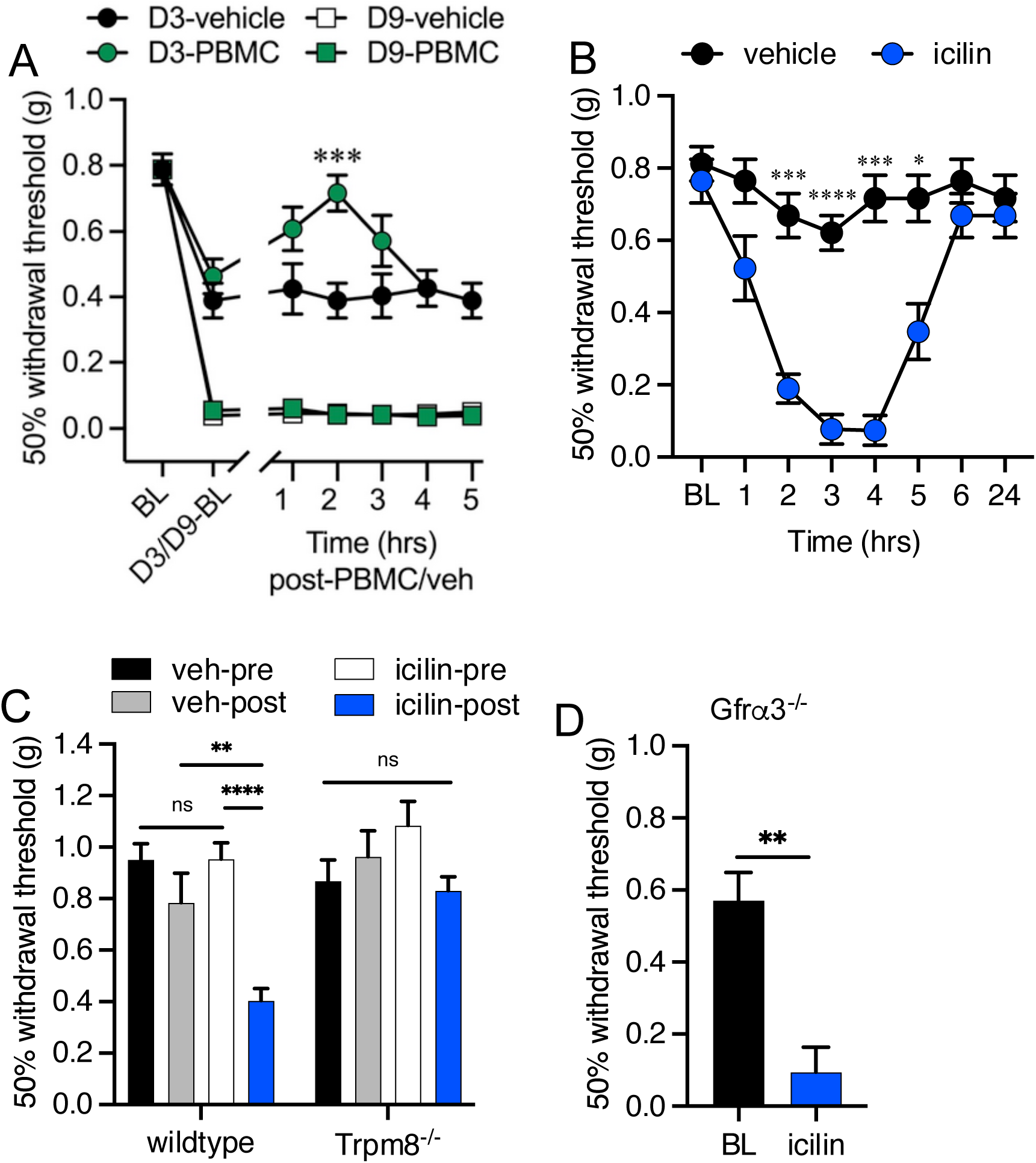
Activation of TRPM8 afferents in the mouse dura induces tactile allodynia. (A) Wildtype mice treated with 1 mg/kg NTG to induce basal periorbital mechanical allodynia at 2 days post-injection (Day 3), as well as after 4 injections repeated every 2 days, measured 9 days after the initial injections, compared to BL, a phenotype that remained in mice subsequently treated with vehicle (n=8, two-way ANOVA with Dunnett’s multiple comparisons test, compared to baseline, p<0.01 on Day 3, p<0.001 on Day 9). However, basal allodynia on Day 3, but not Day 9, was reduced in animals treated with the PBMC at 2 hrs post-injection compared to vehicle treated mice (n=8, two-way ANOVA with Dunnett’s multiple comparisons test, ***p<0.001). (B) Periorbital mechanical sensitivity measured in wildtype mice that received an intradural injection of either icilin or vehicle. Mechanical allodynia was evident by 2 hrs post-injection and lasted for up to 5 hrs (n=6 for each condition, two-way ANOVA with a Bonferroni’s multiple comparisons test for vehicle vs. icilin at each time point; *p<0.05, **p<0.01, ***p<0.001, ****p<0.0001). (C) TRPM8-null (Trpm8^-/-^) mice displayed normal mechanical sensitivity in their hindpaws 3 hrs after an intradural injection of icilin, in contrast to wildtype animals (n=10 for each condition, one-way ANOVA with a Tukey’s multiple comparisons test for vehicle vs. icilin in each condition; ^ns^p>0.05, **p<0.01, ****p<0.0001). (D) Intradural icilin induces periorbital tactile allodynia in Gfrα3^-/-^ mice (3 hrs post-injection, n=8, paired two-tailed t-test, **p<0.01).

These data, and evidence that mouse meningeal afferents express TRPM8 (Fig. 1), suggest that sensitization of TRPM8 channels by artemin and GFRα3 signaling in this tissue leads to migraine-like phenotypes in the mouse, but it is unclear if these are functional channels. Thus, we next asked if direct application of the potent and selective TRPM8 agonist icilin ^9, 57^ to the mouse dura could induce behaviors consistent with mouse migraine-like phenotypes. Previously, Burgos-Vega et al. (2016) demonstrated that supradural application of icilin in the rat induced both cephalic and extracephalic mechanical allodynia that was inhibited by co-application of sumatriptan and L-NAME ^26^. While this effect was sensitive to TRPM8 antagonism, the genetic dependence of this icilin phenotype in mice has not been established. Thus, we directly applied icilin (1 nmol in 5 μl) to the mouse dura and measured changes in periorbital mechanical sensitivity pre- and post-injection, observing robust allodynia that was significantly different than vehicle injected mice by 2 hours, lasting up to 5 hours before returning to baseline (BL) by 6 hours post-injection (Fig. 6B). This time course is consistent with previous findings in the rat ^26^. We also tested extracephalic sensitivity in the mouse hindpaw, observing mechanical allodynia at 3 hours post-injection, sensitization that was absent in Trpm8^-/-^ mice (Fig. 6C). Lastly, in a final test of this hypothetical pathway, we predict that artemin and GFRα3 act upstream of TRPM8 and thus the dura icilin induced allodynia should occur independent of GFRα3, a hypothesis supported by clear periorbital tactile allodynia in Gfrα3^-/-^ mice treated with supradural icilin (Fig. 6D). Thus, TRPM8 expressed in the mouse meninges is functional and activation of these channels leads to both cephalic and extracephalic mechanical allodynia that is consistent with preclinical rodent migraine models.

## DISCUSSION

TRPM8 is the principal detector of environmental cold in the skin but, like the heat-sensitive channel TRPV1, is also present in afferents innervating tissues not involved in thermosensation ^58, 59^. For example, TRPM8-positive afferents extensively innervate the rodent bladder and contain nociceptor markers such as CGRP at a higher percentage than the overall TRPM8-positive somas per ganglion, consistent with the channel’s role in bladder dysfunction ^60–62^. Similarly, TRPV1 is expressed in visceral and muscle afferents in larger proportions compared to those innervating the skin ^63^. Thus, while present and functional, the thermosensitivity of these receptors is likely not relevant in these tissues, indicating that endogenous agonists or modulators sensitize or activate these channels to assert their physiological roles in regions other than the skin ^64, 65^. For example, agonists like menthol and capsaicin sensitize the temperature sensitivity of thermoTRPs like TRPM8 ^9^ and TRPV1 ^66^, respectively, such that the channels gate at physiological temperatures ^67^. Similarly, several signal transduction cascades induced by inflammation sensitize TRPV1 such that the channel is active at skin temperatures ^68, 69^. Thus, their sensitivity to endogenous and exogenous stimuli likely underlies their potential contribution to migraines ^64^. For TRPM8, artemin is the only known endogenous agent that leads directly to TRPM8-dependent hypersensitivity, a pathway that has only been studied in relation to cutaneous cold sensation ^11^. Here, we provide compelling evidence that this pathway also underlies a mechanism whereby TRPM8 sensitization may lead to migraine-like pain.

Artemin is a ligand of the glial cell line-derived neurotrophic factor (GDNF) family and plays crucial roles in nervous system development, nociception, and the pathogenesis of neurological disorders ^70^. At the transcriptional level, artemin mRNA is expressed in developing nerve roots, Schwann cells, vascular and visceral smooth muscle cells, endothelial cells, skin fibroblasts, and immune cells, indicating its potential release from these sites ^71, 72^. The GDNF Family Receptor Alpha 3 (GFRα3) is a high affinity receptor for artemin and shows a similar elevated expression pattern in sensory ganglia, smooth muscle cells, skin fibroblasts, and endothelial cells ^72^. Notably, both artemin and GFRα3 are expressed in dura afferents, with elevated levels in rodent models of migraine, suggesting a significant role in migraine pathogenesis ^37, 38^. Intriguingly, our previous work has demonstrated that artemin, through GFRα3, induces nociception, including cold hypersensitivity, in vivo ^32–35^. This effect is TRPM8-dependent, with GFRα3 expressed in approximately half of TRPM8 neurons in the trigeminal ganglion (TG), and both inflammatory and neuropathic cold pain are absent in GFRα3-null mice ^32, 34^. Peripheral ARTN neutralization alleviates established inflammatory and neuropathic cold pain, highlighting a potential therapeutic intervention ^32, 34, 53^. These findings suggest similarities in ARTN/GFRα3 signaling in both cold and trigeminovascular pain.

Nitroglycerin (NTG), a nitric oxide (NO) donor, can induce headaches in both migraineurs and non-migraineurs by sensitizing the trigeminovascular system, causing vasodilation of meningeal blood vessels, and promoting the release of proinflammatory factors ^73–75^. In this study, we administered a moderate dose of NTG (1 mg/kg, one-tenth of the commonly used dosage ^45^) to induce a significant tactile allodynic phenotype in the cephalic region of mice, minimizing the risk of overwhelming sensitization. We found that the absence of GFRα3 and the neutralization of artemin via a monoclonal antibody prevented acute migraine-like pain triggered by NTG, suggesting that systemic NTG administration could induce artemin release and interaction with GFRα3 at the meninges. Additionally, a chronic migraine-like phenotype established by repeated NTG treatment was also attenuated by artemin neutralization or the absence of GFRα3, indicating sustained increased artemin levels facilitated baseline hypersensitivity in the rodent model of chronic migraine ^45^. These data compellingly demonstrate that the NTG-induced migraine-like phenotype is highly dependent on artemin release and signaling through GFRα3. This significant correlation is underscored by the observed elevation of artemin and GFRα3 levels in the dura mater of NTG-induced rodent migraine models and trigeminal ganglion cultures ^39^. Artemin, which is abundantly expressed in vascular smooth muscle cells, promotes vasodilation by relaxing these cells, and its level positively correlates with inducible nitric oxide synthase (iNOS) levels ^39^. Consequently, neutralizing artemin could mitigate NTG-induced vasodilation, a primary contributor to nociception in this model. Furthermore, the excessive production of nitric oxide in the NTG model can trigger an inflammatory response and subsequent neurogenic inflammation ^76^. The precise molecular mechanisms linking NTG effects and artemin release remain to be elucidated, and uncovering these relationships is crucial for developing future treatments.

A limitation of the NTG model is the systemic nature of its effects, making it difficult to determine if the phenotype is primarily produced in the meninges. However, we found that direct application of artemin to the mouse dura mater triggers cephalic tactile allodynia, a TRPM8-dependent phenotype. Artemin induced allodynia was also inhibited by concurrent treatment with sumatriptan, further supporting this phenotype to be migraine-like. As shown here and in other reports, TRPM8-positive afferents are functionally expressed in mouse meninges, and activation of this channel in this tissue leads to both cephalic and extracephalic tactile allodynia ^26, 27, 52, 77^. This phenotype is reminiscent of our prior work showing that peripheral nociceptor activation by endogenous proalgesics leads to artemin-dependent cold allodynia ^32^ and that subcutaneous artemin application leads to TRPM8-dependent cold pain ^35^. It is also essential to recognize that artemin might not account for all migraine episodes. For instance, in the inflammatory soup migraine model, deleting the GFRα3 receptor only partially alleviates cephalic tactile allodynia induced by inflammatory factors. Hence, the development of migraine-like pain likely involves multiple molecular pathways, necessitating a therapeutic approach that targets multiple pathogenic pathways for effective treatment.

Our results suggest that activating TRPM8 leads to migraine-like pain, but cooling is also reported to have a short-term effect alleviating headaches, and menthol modestly relieves migraines when applied either intranasally or to the forehead ^28–30, 78–80^. Further, one report suggests TRPM8 has a protective effect and contributes to a faster recovery in the chronic migraine model in males but not females ^81^. TRPM8 is the predominant cold sensor in mammals, mediating innocuous cool and noxious cold sensation both acutely and in pathological settings ^9, 33, 41, 46, 48, 82–84^. Conversely, modest thermal or chemical cooling of the skin inhibits both inflammatory and neuropathic pain ^48, 85–87^. These same stimuli also alleviate acute and chronic itch, airway irritation, and aversion to oral nicotine ^88–91^. These processes are TRPM8-dependent and likely occur via TRPM8 afferent activation of spinal cord dorsal horn inhibitory interneurons, suggesting a similar mechanism in the trigeminocervical complex (TCC) may provide headache relief ^89, 92^. As cephalic and extracephalic cutaneous allodynia with migraines is proposed to result from spinal and supraspinal central sensitization, respectively ^93–96^, stimulation of cephalic cutaneous TRPM8 afferents, which converge with meningeal afferents in the TCC, may provide an inhibitory input that reduces sensitization produced in the meninges. Therefore, in the context of migraine-like pain, there may be a distinction between the activity of TRPM8-expressing afferents in the facial skin and those in the meninges, an intriguing posit that requires further investigation.

How can TRPM8 activation lead to migraine-like pain when its only known role in nociception is related to cold sensation? The prevailing model for the sequence of events leading to migraine suggests that a variety of factors alter cortical excitability that eventually leads to activation of meningeal trigeminal nociceptors which induce sensitization of 2^nd^ order neurons in the brainstem ^97, 98^. Our results suggest that artemin/GFRα3 sensitization of TRPM8 in the meninges is one such pathway that can result in migraine and headaches. Further, when considered in the context of the TRPM8 variants identified in GWAS on migraine ^21, 23^, individuals with elevated TRPM8 expression would be more susceptible to an increase in artemin or other endogenous agents that sensitize TRPM8 channels, perhaps underlying their predisposition to migraines ^21^. This hypothesis requires further study, but our results now implicate artemin/GFRα3/TRPM8 as novel players in the molecular mechanisms of migraine.

## Acknowledgements

The authors wish to thank current and former members of the McKemy Laboratory for their assistance in the completion of this work.

